# Disentangling the mechanisms by which nitrogen enrichment affects consumer damage at different organisational levels

**DOI:** 10.1101/2023.11.24.568521

**Authors:** Tosca Mannall, Caroline Daniel, Vera Alessandrello, Anne Kempel, Eric Allan

**Affiliations:** Institute of Plant Sciences, University of Bern, Altenbergrain 21, 3013 Bern, Switzerland; WSL Institute for Snow and Avalanche Research SLF, Flüelastrasse 11, 7260 Davos, Switzerland; Climate Change, Extremes and Natural Hazards in Alpine Regions Research Center CERC, Flüelastrasse 11, 7260 Davos, Switzerland; Centre for Development and Environment (CDE), University of Bern, Mittelstrasse 43, 3012, Bern, Switzerland

**Keywords:** herbivory, pathogens, global change, nitrogen, biodiversity, PaNDiv Experiment, resource economic strategies

## Abstract

Nitrogen enrichment could affect primary consumers by increasing foliar N content (nitrogen-disease hypothesis), by increasing dominance of palatable, fast-growing plants (growth-defence trade-off) or by reducing plant diversity (resource concentration effect). These mechanisms might operate differently at different organizational levels. We tested this in a grassland experiment (PaNDiv), manipulating nitrogen enrichment, plant species richness, fast-slow functional composition, and foliar pathogens. We assessed herbivory and pathogen damage, on focal plant individuals, species, and communities. Plant community characteristics were more important drivers of consumer damage than nitrogen enrichment. Host concentration effects strongly affected pathogens but mostly at the species level. Growth-defence trade-off effects were widespread but frequently emerged from interactions between species (associational effects) or with resource concentration effects, leading to different patterns at species and community levels. Our findings suggest that key mechanisms may operate differently at different organisational levels calling for better understanding of how consumer effects scale across levels.

## Introduction

Insect herbivores and fungal pathogens are key plant enemies that have large impacts on diversity and ecosystem functioning (Allan *et al*. 2010; Mordecai 2011), and that are strongly affected by global change (Burdon *et al*. 2006; van Klink *et al*. 2020). It is critical to understand the mechanisms behind effects of global change, such as nitrogen enrichment, on consumer groups. In addition to increasing tissue nutrient levels, nitrogen enrichment typically causes a shift from slow growing, conservative plants to fast growing acquisitive species (Suding *et al*. 2005; Lavorel & Grigulis 2012; Lind *et al*. 2013; Borer *et al*. 2014), and a loss of plant species richness through an increase in light competition (Hautier *et al*. 2009; Liu *et al*. 2016; Eskelinen *et al*. 2022). Nitrogen enrichment could therefore affect consumers via three main mechanisms: 1) a shift in plant nitrogen content and biochemistry, potentially leading to an increase in consumer feeding (nitrogen disease hypothesis, (Huber & Watson 1974; Strengbom & Reich 2006; Dordas 2008), 2) a shift in plant functional composition towards faster growing and poorly defended plants, leading to an increase in consumer attack (growth-defence trade-off hypothesis, (Coley *et al*. 1985; Herms & Mattson 1992), and 3) declining plant diversity leading to increased attack by specialist consumers (resource concentration effect; (Root 1973). The relative importance of these different mechanisms may depend on the group of consumers, and the organisational level at which they are tested.

It may not be straightforward to scale consumer impacts across organisational levels, from individual plants in neighbourhoods, to individual species in patches, to whole communities of several plants species (Table 1). Interactions between individual plants or species may lead to different effects of consumers at different levels. For instance, at the community level, damage may be higher than expected from the individual species present if attractive species promote spillovers and increased damage for co-occurring species, or it may be lower than expected if defended species provide associational resistance to the community. Different effects may also occur at the individual plant and species scale, depending on the consumer group. Fungal pathogens may respond more strongly to small scale (a few 10cms) variation in neighbourhood properties (Barbosa *et al*. 2009; Liu *et al*. 2023), meaning that effects of the surrounding neighbourhood may be stronger for individual plants than for species in larger patches. Most insect herbivores actively search for their host plants and may respond to variation in plant neighbourhoods only at larger scales. In addition, insect herbivores may also be able to better select their host plants, leading to more variation in insect herbivore attack between plant individuals and species in a community, which might make it harder to scale effects from individuals to species and communities. Some studies have looked at different effects of consumers at different spatial scales (e.g., Liu et al. 2023) but few have compared effects between organisational levels.

**Table 1:**
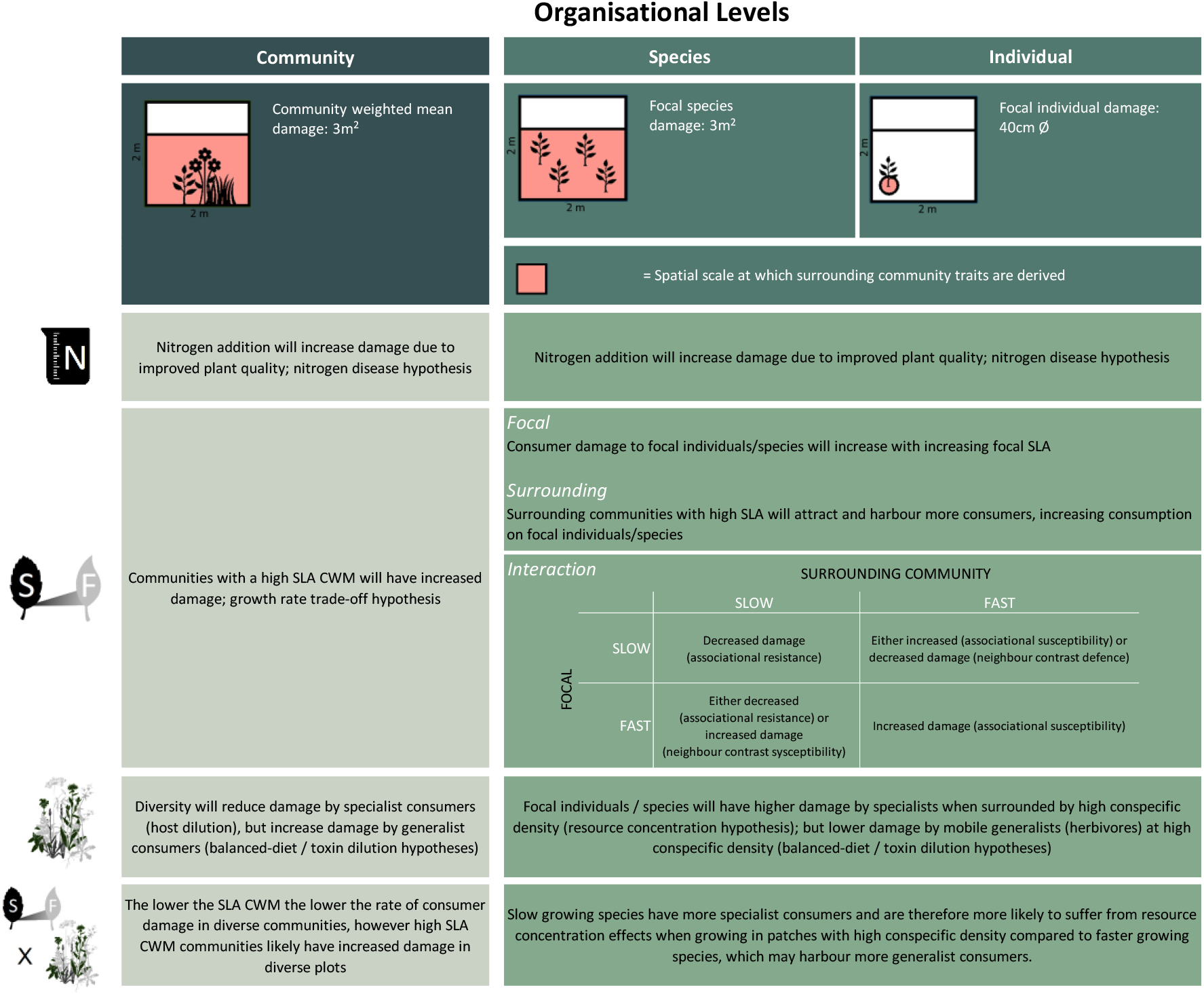
Hypotheses determining how nitrogen can indirectly influence how consumers interact with plant community traits at different organisational levels and scales to influence consumer damage to focal individuals, populations, and plant communities as a whole. Figures from top to bottom: Nitrogen enrichment, Functional composition (fast vs slow plant communities indicated by Specific Leaf Area (SLA)), plant species richness and interaction between functional composition and plant species richness.

Changes in foliar nitrogen content are likely to have a large effect on phytophagous consumers and their impact (Throop & Lerdau 2004; Ebeling *et al*. 2021). Increasing leaf nitrogen concentrations can increase foliar fungal disease severity (nitrogen disease hypothesis, Huber & Watson 1974; Strengbom & Reich 2006; Dordas 2008) and insect survival, development, and reproductive rates (Herms & Mattson 1992; Throop & Lerdau 2004). Nitrogen enrichment is therefore expected to increase consumer damage and its effects are expected to be similar across organisational levels.

A shift towards faster growing plants may also strongly affect consumers. Fast growing plants, with high Specific Leaf Area (SLA, Wright *et al*. 2004), are generally more attractive to consumers because they allocate resources towards growth at the cost of defence (growth defence trade-off hypothesis, Coley et al.1985; Herms & Mattson 1992) and typically have high leaf nutrient contents (Díaz *et al*. 2016). In addition, the traits of neighbouring plants may affect consumer feeding on a focal individual, potentially leading to an interaction between the traits of the focal and those of the neighbours. For example, more palatable and susceptible neighbours may attract consumers to a patch, resulting in increased consumption of the focal individual (associational susceptibility), or spillovers to it (Thomas 1986; Hjalten *et al*. 1993), while well defended and less attractive neighbours may provide protection for a focal individual, reducing its damage (associational resistance; (Tahvanainen & Root 1972; Atsatt & O’Dowd 1976; Alm Bergvall *et al*. 2006). However, opposing patterns are also possible (see Table 1). Associational effects (Barbosa et al. 2009) could lead to different effects of consumer at the species and community level, e.g., fast growing plant species and communities dominated by fast growing plants may not receive similarly high attack. Consumer attack could be amplified in communities with only palatable (fast growing) plants, or it could be reduced in communities with a few defended (slow growing) species, which reduce consumer damage below that expected based on the mean growth strategy of plants in the community.

In addition, plant diversity can strongly impact consumer feeding, although its effects may vary between consumer groups. The resource concentration hypothesis (Root 1973) predicts that host specific consumers (such as many sap sucking Hemiptera, Forister et al., 2015) are likely to locate and remain in stands with high host density. Conversely, generalist herbivores (e.g., many Orthopterans; Forister et al., 2015) might profit from a more balanced diet and perform better in diverse patches (Pfisterer *et al*. 2003). Foliar fungal pathogens, which are mostly dispersed randomly, are impacted by changes in the relative abundance of susceptible versus resistant hosts (Liu *et al*. 2017; Halliday *et al*. 2019). For specialist pathogens (such as rusts; Rottstock *et al*. 2014b), infection rates are also likely to increase with conspecific density (host dilution effects; (Mitchell *et al*. 2003; Rottstock *et al*. 2014) as spore transmission is easier in monospecific stands (Alexander & Jeanne 2000; Lopes *et al*. 2016). However, generalist pathogens (such as those causing leaf spots) might spill over onto a focal individual surrounded by susceptible, heterospecific neighbours (Power & Mitchell 2004; Parker *et al*. 2015). Generalist vector-transmitted pathogens may also spread more easily in diverse host communities (Power & Mitchell 2004; Halliday *et al*. 2017). Fungal pathogen reduction, using fungicides, could alter the strength of resource concentration effects because fungicides do not kill all pathogenic fungi and could affect specialist and generalist groups differently (Cappelli *et al*. 2020). We might expect that resource concentration effects at the population or species level do not always lead to negative effects of plant diversity on consumer attack at the community level, if spillovers of generalist pathogens and dilution of specialists occur simultaneously in diverse plant communities.

These mechanisms are likely to interact to influence plant-consumer interactions. Slow growing species may have more specialist consumers, who can detoxify their chemical defences (Krieger *et al*. 1971) or feed on leaves with high structural defences, and might be more likely to suffer from resource concentration effects. Interactions may also occur between consumer groups. For example the performance of insect herbivores can be increased by fungal pathogens, as pathogens serve as a nutritious resource, particularly for chewing insect herbivores (Eberl *et al*. 2020), or because defences against pathogens and insects trade off (Felton & Korth 2000; Stout *et al*. 2006; Thaler *et al*. 2012). However, insect performance can also be decreased by fungal pathogens (Jallow *et al*. 2004) as pathogen infection could prime plants to increase overall defences (Conrath *et al*. 2002; Ali & Agrawal 2012). Hence, the presence of fungal pathogens may alter the strength of resource concentration or growth-defence effects for insects, and this might differ between herbivore guilds.

Here we test how nitrogen enrichment indirectly influences leaf damage by four guilds of invertebrate herbivores and foliar fungal pathogens. Using a grassland experiment factorially manipulating nitrogen, plant species richness, fast-slow plant functional composition (mean specific leaf area) and fungicide, we tested the relative importance of growth-defence trade-off, host concentration and direct nitrogen effects, in influencing consumer damage at different organisational levels and corresponding spatial scales. We calculated damage on planted *individuals* (phytometers), focal *species* and *communities* and tested the effect of focal characteristics at each level. We also looked at how characteristics of the surrounding community affected damage at the individual (surrounding community of 40 cm ∅) and species (surrounding community of 3m^2^) levels.

We asked the following questions:

I. How does nitrogen addition affect consumer damage at different organisational levels?
II. How does a shift towards fast growing (high SLA) plants affect consumer damage at different organisational levels?
III. How does a reduction in plant diversity affect consumer damage at different organisational levels?
IV. How do these effects interact to influence consumer damage at different organisational levels?

## Methods

### PaNDiv field experiment

The PaNDiv field experiment was established in October 2015, in Münchenbuchsee (Bern, Switzerland, 47°03′N, 7°46′E, 564 m a.s.l.) and factorially manipulates plant species richness, plant functional composition, nitrogen enrichment and fungal pathogen removal (Pichon *et al*. 2020). The experiment comprises 216 2 x 2m plots, separated by one metre paths, arranged in four blocks. A 0.5 x 2m strip was left unweeded on each plot, meaning the weeded area of the plot is 3m^2^ (Figure S1).

The species pool consisted of 20 grass and herb species typical of mesic, central European grasslands (Table S1). Species were divided into two pools corresponding to different resource economic strategies (Wright *et al*. 2004), using leaf nitrogen content and SLA. The ‘Fast’ species pool included ten fast-growing, acquisitive species with high SLA and leaf nitrogen. The ‘Slow’ species pool comprised ten species with a more conservative strategy, and low SLA and leaf nitrogen.

The plant diversity treatment included four diversity levels: 1, 4, 8, and 20 species mixtures. Species diversity levels are maintained by removing non-target species by weeding, which occurs three times per year (April, July and September). Plots with four and eight species contain either only fast or only slow growing species, or a combination of the two strategies. Species compositions for each plot were randomly selected from the respective species pool, with the constraint that all plots (except monocultures) had to contain both grasses and herbs. Each unique species composition received the four combinations of nitrogen enrichment and fungicide application.

Nitrogen treated plots received urea twice a year (beginning of April and end of June), resulting in an annual addition of 100 kg N ha^-1^y^-1^, which corresponds to intermediately intensive grassland management (Blüthgen *et al*. 2012).

Foliar fungal pathogens were suppressed using two fungicides (“Score Profi”, 24.8 % Difenoconazol 250 g.L-1 and “Ortiva”, 32.8% Chlorothalonil 400 g.L-1 6.56% Azoxystrobin 80 g.L-1), applied four times during the growing season (beginning of April and June, late July and September). Control plots were sprayed with the same amount of water. The diversity gradient was crossed with fungicide and nitrogen treatments, resulting in 80 monocultures, 60 four, 60 eight and 16 twenty species plots. The entire experimental field site was mown twice per year, in mid-June and mid-August, following intermediate-extensive grassland management in the region. For further details on the PaNDiv Experiment see Pichon *et al*. (2020).

### Individual plants

Phytometer individuals were planted within the PaNDiv plots in September 2019, into neighbourhoods varying in the abundance of conspecific and heterospecific neighbours. We defined the neighbourhood as all plants within a circle of 40cm diameter around the focal phytometer (Figure S1). Phytometers were planted into monocultures and mixture plots. For the mixture plots we largely used a set of 120 four and eight species plots that had been part of the original experimental design but where weeding had ceased after 2018 (the nitrogen and fungicide treatments continued to be applied). We avoided neighbourhoods in these plots with a large number of non-target species (i.e., species not in experimental species pool). Phytometers comprised 10 out of the 20 PaNDiv experimental species (Table S1) and had between 40 and 55 individuals per species per treatment.

### Plant species and community characteristics

We measured specific leaf area (SLA = surface area/dry weight, *m^2^ kg^-1^*) in June 2019, by collecting one leaf from five individuals in each of the 80 monoculture plots. We rehydrated the leaves in the lab and measured leaf surface area and dry weight according to Garnier et al. (2001). The SLA per species and treatment is the mean across the five samples. These SLA values were used to calculate the **SLA of the focal individual and species**, meaning that we incorporate intraspecific trait variation in response to N addition and fungicide but not in response to the different community contexts.

To characterise the **SLA of the surrounding community**, we visually estimated the percentage cover of the surrounding species at the different organizational levels. At the “species” level, we recorded the cover of each of the target plant species (PaNDiv experimental species) in the central 1m^2^ (Figure S1) of each plot, between 25^th^ May and 5^th^ June 2020. Percentage cover was converted into relative cover, by dividing the percentage cover of each species by the total percentage cover. At the “individual” level, we assessed the percentage cover of each species in a 40 cm diameter circle surrounding each phytometer in June 2020.

We then calculated the surrounding plant community **SLA CWM**. These were calculated as CWM values for SLA as:

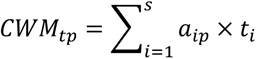

where *a_ip_* is the relative percentage cover of species *i* in plot *p*(3m^2^ plot for species level, 40 cm diameter plot for the individual plant level) and *ti* is the mean trait value of species *i* in the monoculture of the same treatment (i.e., control or nitrogen/fungicide addition). We included conspecifics in the calculation of the surrounding CWM values because many (300) phytometers were planted in monocultures where they only had conspecific neighbours and removing these would have dramatically reduced the gradient in conspecific density. The plot CWM SLA was used as the focal SLA measure for the community level.

**Conspecific density** was then calculated as the relative percentage cover of the focal species, in a 3m^2^ or 40 cm diameter plot. At the community level we used the diversity of the plot to test for resource concentration effects. In line with standard approaches in biodiversity-functioning research (Schmid *et al*. 2002), we used the sown diversity of the plot.

### Consumer damage

#### Species and community levels

Insect herbivory was visually assessed from the 25^th^ May – 8^th^ of June 2020 on all 216 plots. Five individuals of each target species were haphazardly selected from the central 1 m^2^ of each plot (Core, Figure S1) and were only supplemented by individuals from the remainder of the weeded plot when insufficient individuals were found in the centre. Damage was characterised by the type of herbivory; chewing, sucking, galls and leaf mining, following (Loranger *et al*. 2014). The percentage of each damage type was recorded on each individual, excluding juvenile and senescent leaves.

Overall infection incidence of foliar fungal pathogen guilds (rusts, powdery mildews, downy mildews and leaf spots, see Rottstock *et al*. (2014b), was assessed for each plant species in each plot from the 8^th^ - 14^th^ October 2020. We haphazardly selected 10 individuals per species, from the core of each plot (**Error! Reference source not found.**) and recorded the proportion of infected individuals. If there were < 10 individuals, infection incidence was calculated from all individuals present. Fungal infection was measured as incidence, firstly, because it was positively correlated with infection severity (Cappelli *et al*. 2020), and secondly because fungal infection is partially internal and cannot be measured in its entirety via visual inspection. Herbivory on the other hand, is defined here as the removal of leaf tissue, and is therefore more easily identifiable externally. To assess the overall community consumer damage per plot, we calculated the community weighted mean of each damage guild, by multiplying the damage per plant species with the percentage cover of that species on a given plot.

#### Individual level

Damage by herbivores and pathogens was assessed in June 2020 on each phytometer and was measured as the percentage of 5 leaves damaged by each herbivore and pathogen guild (as above). A total of 1860 phytometers (1666 after accounting for mortality: with between 108 - 191 individuals of each phytometer species) were assessed for damage.

### Statistical analyses

We analysed variation in chewing and sucking herbivory, and rust and leafspot foliar fungal infection, at three organizational levels. At the **Individual level** by assessing the response of focal individual consumer damage, to community characteristics from the surrounding **40cm⌀ neighbourhood**, at the **species level** by assessing the response of plot level focal species consumer damage, to community characteristics from the surrounding **3m^2^ plot**, and at the **Community level** by assessing the response of plot level community weighted mean (CWM) consumer damage, to community characteristics of the **3m^2^ plot**. Note that the nitrogen and fungicide treatments are applied at the plot scale and therefore do not vary between neighbourhoods within a plot. We excluded gall and mining herbivory and mildew pathogen infection because these damage types were rare and restricted to particular plant species. All selected consumer damage guilds were modelled separately at all levels using the function *glmmTMB*, R package *glmmTMB* (Brooks *et al*. 2017), as it allows the flexible fitting of error distributions. All analyses were performed in R version 4.1.2 (R Core Team 2021).

At the individual and species level, the full model for each consumer response variable included the following fixed effects: nitrogen(N), fungicide(F), conspecific density, surrounding species community weighted mean SLA (surrounding SLA), focal SLA and two-way interactions between variables:

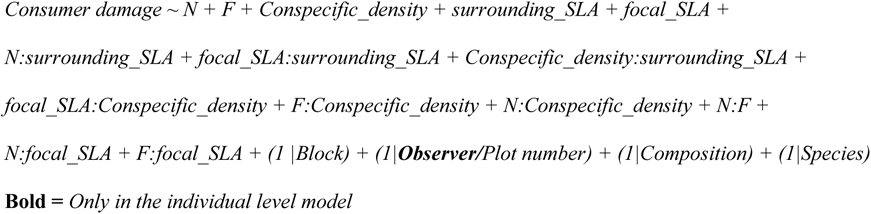

For the community level models, plot species richness replaced the conspecific density measure and focal SLA was replaced by CWM SLA, surrounding SLA was not included. Random effects were included in all models to account for the experimental setup: block (4 levels), plot number (216 levels), composition (51 levels), and focal plant species (20 or 10 levels), with the addition of observer (12 levels) in the individual level models to account for multiple observers who assessed damage and percentage cover.

For each response variable, the best fitting distribution family was selected for each glmmTMB model based on the distribution of the residuals (simulateResiduals function in DHARMa package version 0.4.4, Hartig & Lohse, 2021) and AIC when multiple distributions had adequate residual distributions. Resulting in the community scale leafspot model having a gaussian distribution, and the remainder of models having Tweedie distributions (Dunn 2021). Effects plots and results tables were created using the plot_models and tab_model functions respectively from the sjPlot package version 2.8.10 (Lüdecke 2021). All other plots used predicted values derived from the glmmTMB models, using ggpredict from ggeffects package version 1.1.1 (Lüdecke 2018) and were plotted using ggplot from ggplot2 package version 3.3.5 (Wickham 2016).

## Results

Overall, we observed that shifts in the functional composition and diversity of the plant community had a greater influence on consumer damage than the nitrogen treatment. The nitrogen treatment only increased chewing herbivory on phytometers at the individual level (Figure 1, Table S2, P=0.001). There was no other significant effect of nitrogen either alone, or in interaction with other factors, at any level, for any consumer guild.

**Figure 2:**
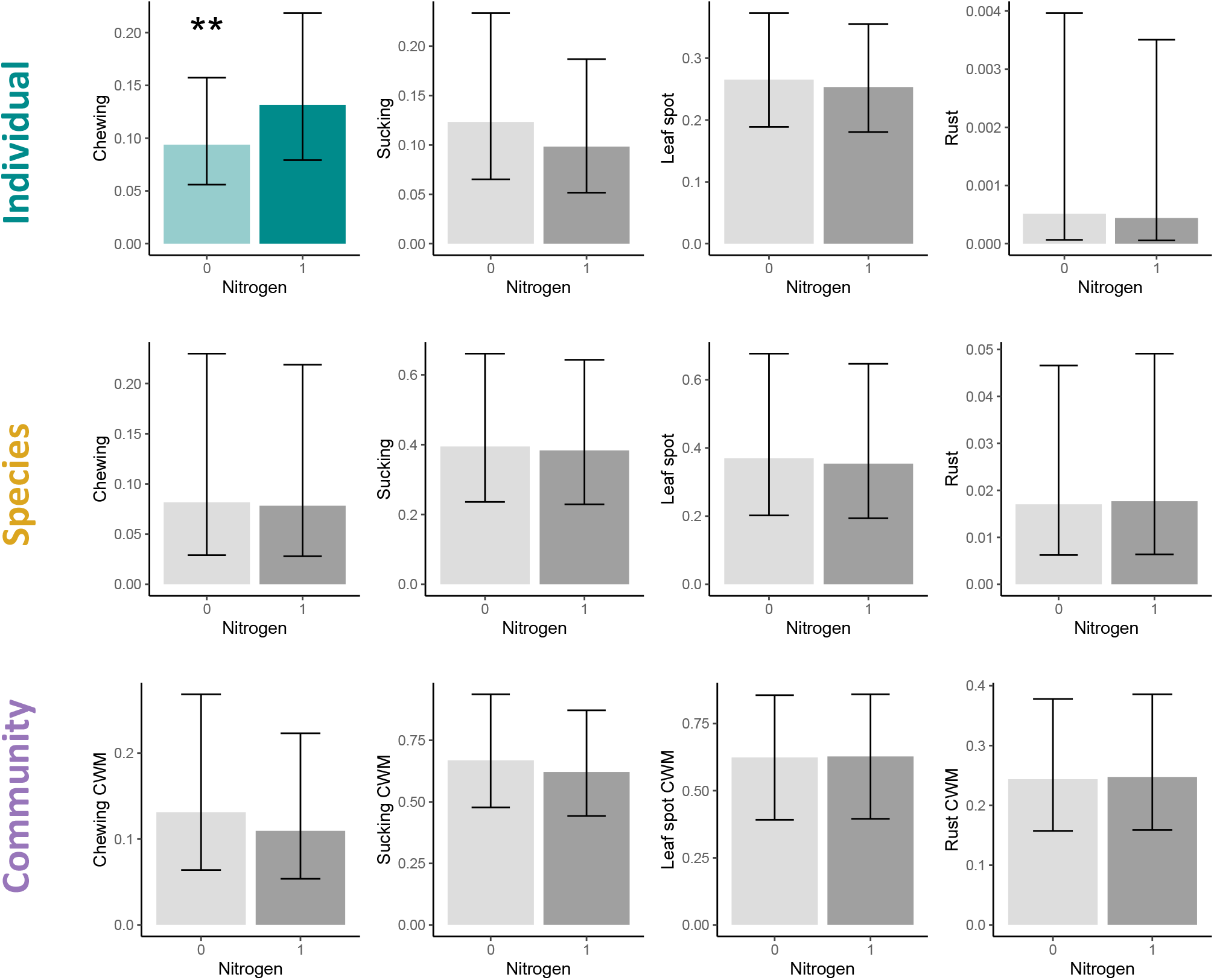
Effect of nitrogen treatment (100 kg N ha-1y-1) on consumer damage at three different levels. Means with 95% confidence intervals, predicted from glmmTMB models are shown. Top row: Individual level(40cm⌀), consumer damage measured as the percentage of leaves damaged on the focal individual; Middle row: Species level=spatial scale 3m^2^, insect herbivory measured as mean percentage of leaves damaged/plant/species/plot, and pathogen infection measured as the proportion of plants infected/species/plot; Bottom row: Community level = spatial scale 3m^2^, consumer damage measured as community weighted mean (CWM) percentage of leaves damaged/plant/species/plot (insect herbivory), or as the proportion of plants infected/species/plot (pathogen infection).

We found several effects of the SLA of the focal or surrounding plants, often in interaction and mostly at the species and community levels. However, the focal SLA (of individual plants or species) alone had minimal effects on consumer damage. It only influenced chewing herbivory at the individual level, where increasing phytometer SLA decreased chewing damage (Figure 2, Table S2, P=0.035). Focal SLA had larger effects at the community level where it increased infection by rust fungi (Table S3, P=0.002). Fast growing plant species experienced the most rust infection when growing in fast growing communities, while slow species did not suffer more rust infection when surrounded by fast species (Figure 5A, Table S4, P=0.029). The surrounding SLA also affected chewing damage, at the species level. As we hypothesised, chewing herbivory increased on species growing in plots with high surrounding SLA (Figure 3, Table S2, P=0.007), but this effect was predominantly driven by slow growing focal species being damaged more when growing in plots surrounded by fast species (Figure 5 B, Table S2, P=0.004). Surrounding SLA had no effect on leaf spots at any level.

**Figure 3:**
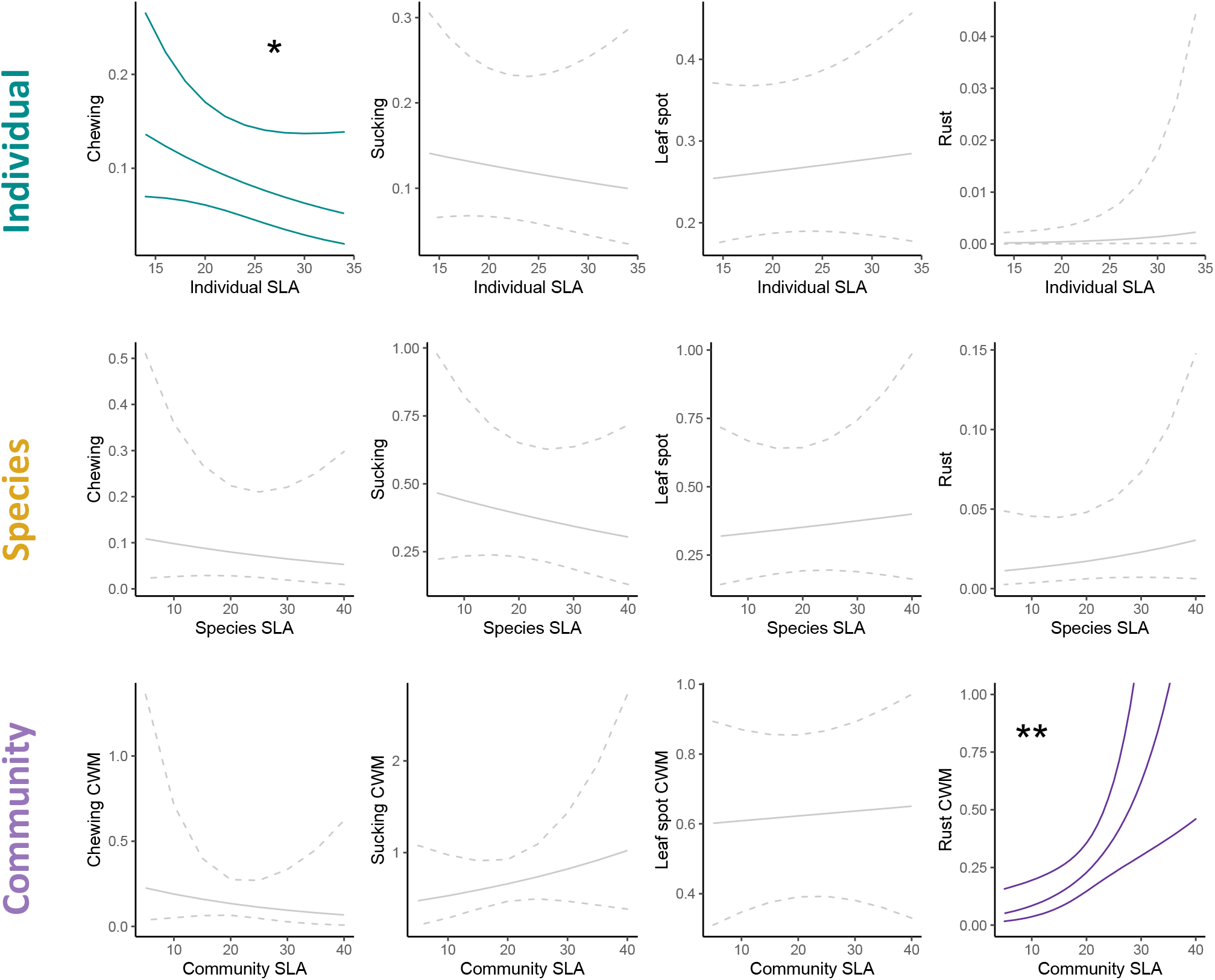
Effect of focal individual/species/community specific leaf area (SLA) on consumer damage at three different organisational levels. Regression line with 95% confidence intervals, predicted from glmmTMB models are shown. Top row: Individual level = spatial scale 40cm⌀, consumer damage measured as the percentage of leaves damaged on the focal individual; Middle row: Species level =spatial scale 3m^2^, insect herbivory measured as mean percentage of leaves damaged/plant/species/plot, and pathogen infection measured as the proportion of plants infected/species/plot. Bottom row: Community level = spatial scale 3m^2^, consumer damage measured as community weighted mean (CWM) percentage of leaves damaged/plant/species/plot (insect herbivory), or as the proportion of plants infected/species/plot (pathogen infection).

**Figure 4:**
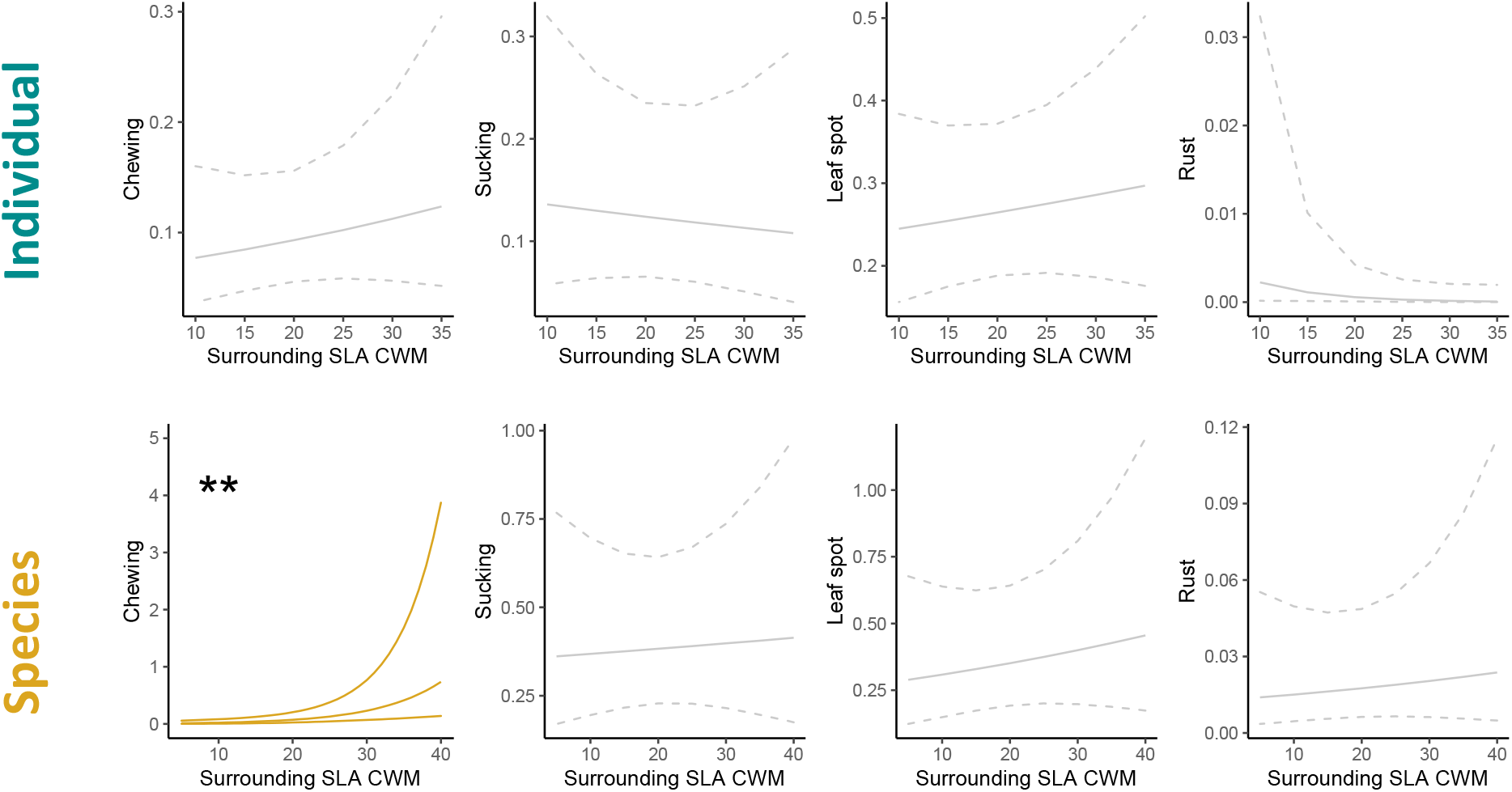
Effect of the community weighted mean (CWM) of specific leaf area (SLA) in the surrounding community, on consumer damage at three different organisational levels. Regression line with 95% confidence intervals, predicted from glmmTMB models are shown. Top row: Individual Level = spatial scale 40cm⌀ of surrounding community, consumer damage measured as the percentage of leaves damaged on the focal individual; Bottom row: Species level= spatial scale 3m^2^ of surrounding community, insect herbivory measured as mean percentage of leaves damaged/plant/species/plot, and pathogen infection measured as the proportion of plants infected/species/plot.

Resource concentration effects were the strongest drivers of pathogen infection and increased infection at both the individual and species level. Interactions between the SLA of the focal species and conspecific density were also important (Figure 5). Increasing conspecific density increased leafspot infection at both the individual (Figure 4, Table S4, P <0.001) and species level (Figure 4, Table S4, P =0.002). Conspecific density also increased rust infection at the species level (Figure 4, Table S4, P <0.001), which was predominantly driven by slow species being more susceptible to infection when growing in high conspecific density patches (Figure 6, Table S4, P= 0.006). Sucking herbivory was not as strongly influenced by conspecific density as predicted but did significantly increase with increasing conspecific density at the species level, for fast growing species (Figure 6 D, Table S2, P=0.042). Chewing herbivores consumed more in diverse communities. Chewing herbivory declined with increasing conspecific density at the individual level (Figure 4, Table S2, p=0.037) and increased with increasing plant species richness at the community level (Figure 4, Table S3, P=0.018). Interactions between consumer guilds were also important. Chewing herbivores caused more damage in species surrounded by high conspecific density, but only when fungicide had been applied, suggesting interactive effects between fungal pathogens and chewing herbivores (Figure S2, Table S2, P=0.023).

**Figure 5:**
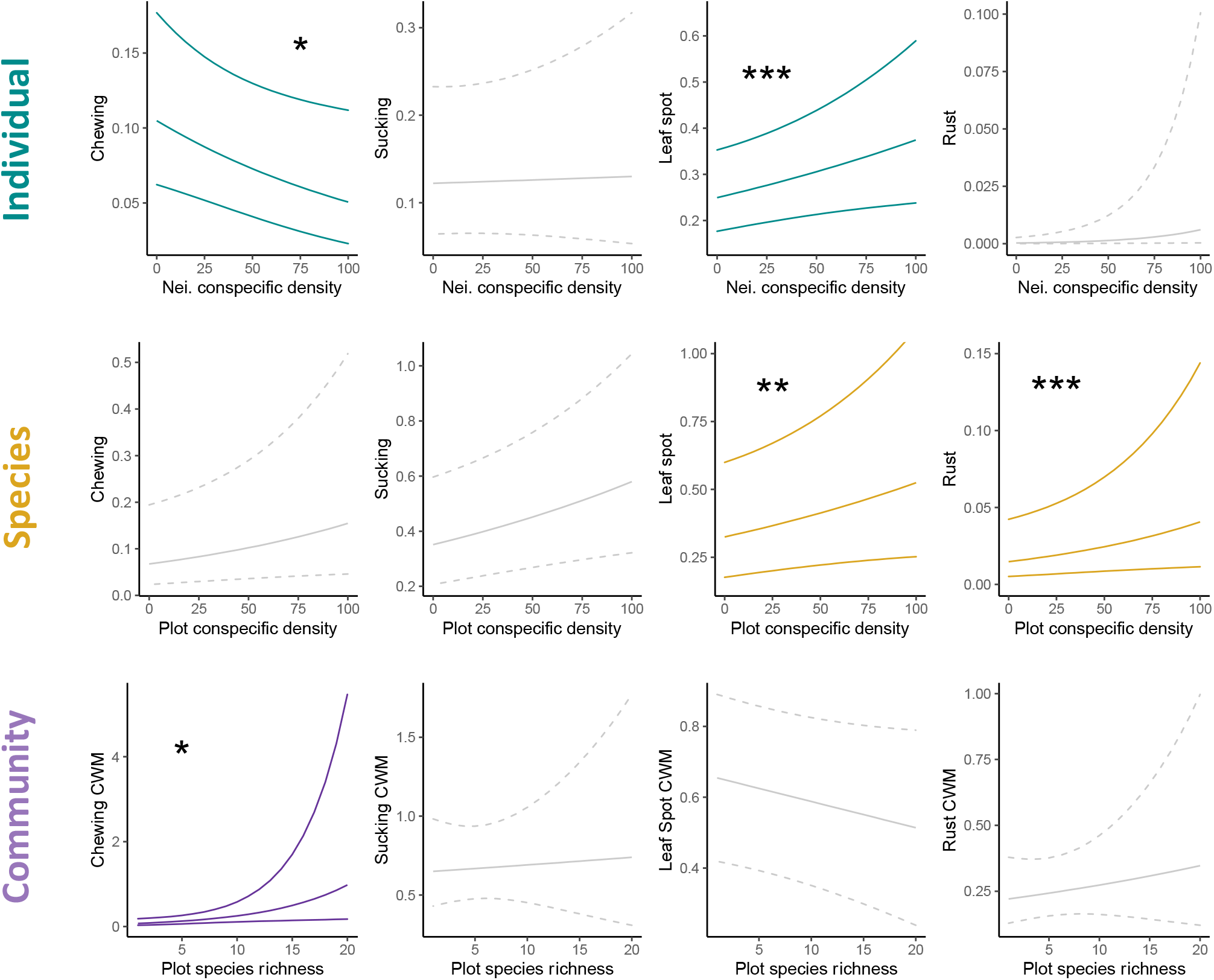
Effect of diversity (conspecific density), on consumer damage at three different organisational levels. Regression line with 95% confidence intervals, predicted from glmmTMB models are shown. Top row: Individual level = spatial scale 40cm⌀, consumer damage measured as the percentage of leaves damaged on the focal individual; Middle row: Species level= spatial scale 3m^2^, insect herbivory measured as mean percentage of leaves damaged/plant/species/plot, and pathogen infection measured as the proportion of plants infected/species/plot; Bottom row: Community level = spatial scale 3m^2^, consumer damage measured as community weighted mean (CWM) percentage of leaves damaged/plant/species/plot (insect herbivory), or as percentage of plants damaged/species/plot (pathogen infection).

**Figure 5:**
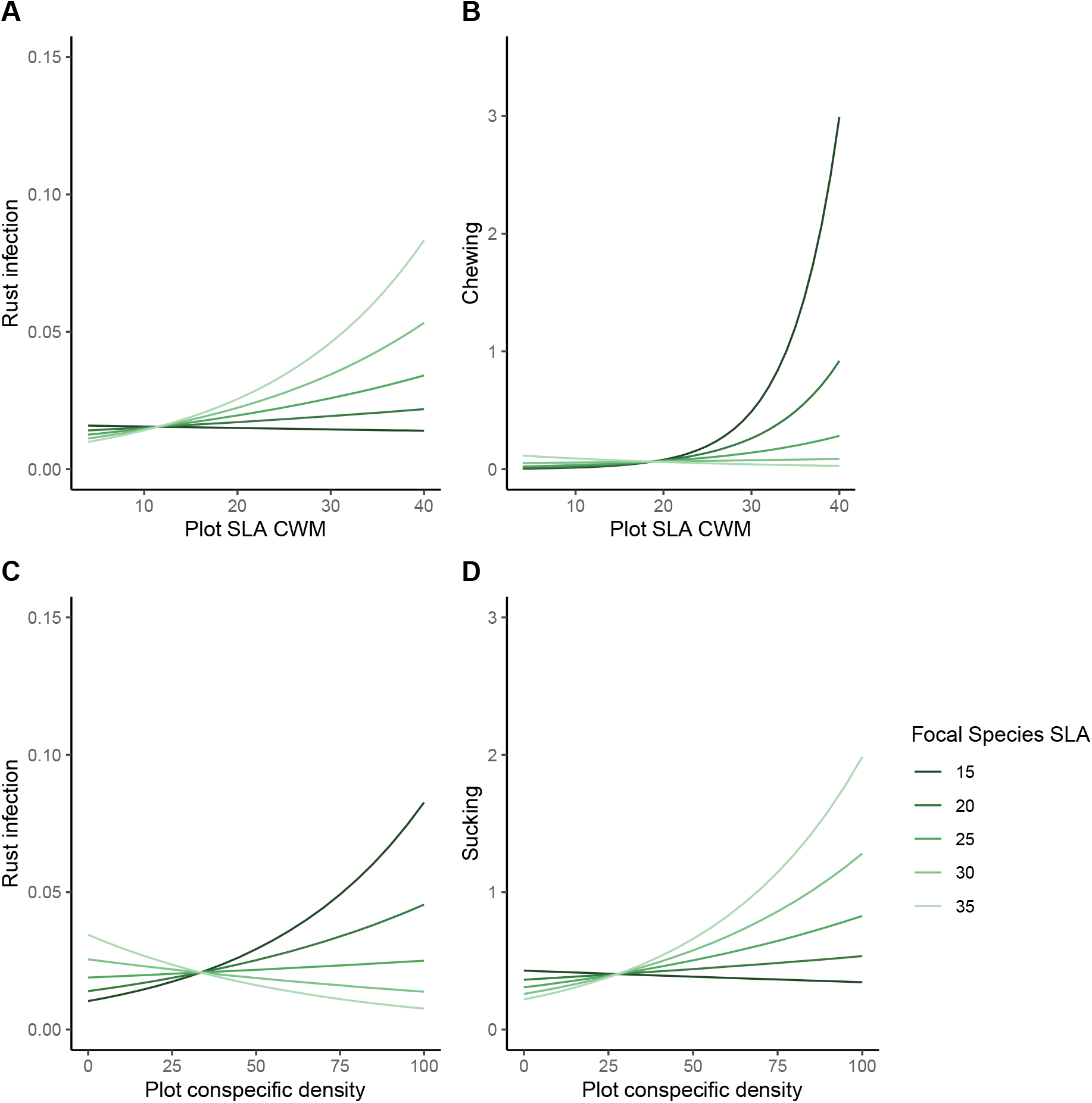
Effect of the interaction between plot level plant community characteristics and focal species specific leaf area (SLA), on consumer damage. Regression lines predicted from glmmTMB models are shown. Spatial scale= 3m^2^, insect herbivory measured as mean percentage of leaves damaged/plant/species/plot, and pathogen infection measured as the proportion of plants infected/species/plot. A) and B), interaction between focal species SLA and plot SLA community weighted mean (CWM). C) and D), interaction between focal species SLA and plot conspecific density.

## Discussion

Overall, we found that plant community composition and diversity had a greater influence on consumer damage than nitrogen fertilisation, supporting previous results from our experiment (Cappelli et al. 2020). We found strong support for the growth-defence trade-off hypothesis as a driver of consumer damage, however, effects varied between consumer groups and depended on the traits of both focal and neighbouring species. Resource concentration effects were also important and were the strongest driver of fungal pathogen infection. However, both growth-defence and resource concentration effects varied in their importance between levels of organisation, often appearing only at the species or at the community level. This occurred because of interactions between species within communities which either dampened or amplified effects from the species to community level. These results suggest that nitrogen enrichment would mostly indirectly affect consumer damage and would do so through a range of different mechanisms at different levels.

The nitrogen treatment only had minimal effects on herbivore damage, suggesting that increasing foliar nitrogen generally did not increase consumer damage. The only significant effect of nitrogen was on chewing herbivory at the individual level. It is surprising to only see the effect at the individual level because we would expect that all plants would be equally attractive on a fertilised plot. However, the phytometers were planted as seedlings, which usually have more palatable and less defended leaves (Fenner *et al*. 1999), and the nitrogen addition might have made them even more palatable to mobile consumers such as orthopterans, lepidoptera larvae or gastropods (Hanley *et al*. 1995). The absence of significant effects of nitrogen on any other consumer guild at any level, provides no evidence for the nitrogen disease hypothesis (Huber & Watson 1974; Mitchell *et al*. 2003; Strengbom & Reich 2006; Dordas 2008). The lack of nitrogen effects could have arisen through a balance between decreasing consumption in high-resource quality (fertilised) patches (Slansky & Feeny 1977; Berner *et al*. 2005; Behmer & Joern 2008) and an aggregation of insect herbivores in fertilised patches (Throop & Lerdau 2004; La Pierre & Smith 2016). Alternatively, nitrogen addition could cause plants to upregulate their defences, thereby reducing consumer damage (Mur *et al*. 2017). Our results differ from studies showing that nitrogen addition increases herbivory (Ebeling *et al*. 2021), however previous studies did not control for indirect effects on nitrogen on plant community composition and these compositional changes may have been the driver of increased herbivory following fertilisation.

Growth-defence trade-off effects were observed for both herbivores and pathogens, but they often depended on the surrounding community. Chewing herbivores were strongly affected by focal individual specific leaf area (SLA), surrounding SLA and their interaction. The reduced chewing damage on focal individuals with high SLA is surprising. However, at the species scale we found that species were most attacked by chewing herbivores in fast growing communities, and slow species were particularly susceptible when growing in fast communities, which is consistent with associational susceptibility effects (Champagne *et al*. 2016). This interaction was weaker (and non-significant) at the individual level but may still have contributed to increased chewing herbivory on slow species. In contrast, fast growing plants experienced the most rust infection when in fast growing communities, while slow species did not suffer infection even in fast communities, suggesting that rusts principally attack fast plant species and do not spillover from fast to slow growing species. At the community level this led to a strong increase in overall pathogen infection with SLA, consistent with the growth-defence trade-off as a main driver of differences in rust infection between communities (Cappelli *et al*. 2020). However, at the species level, fast plants did not suffer more infection than slow species because they gained protection when growing in slow communities (associational resistance). Both the traits of the focal individuals and the traits of the surrounding community were therefore important and led to a strong amplification of rust infection in fast communities. We find evidence for both associational resistance and susceptibility depending on the consumer group. These interactions make it complex to upscale effects of plant traits on consumer feeding from species to communities and highlight the importance of comparing effects between organisational levels.

An increase in conspecific density consistently increased pathogen infection at both the individual and species level. The strong increase of both leafspot and rust infection with increasing host density provides support for the resource concentration hypothesis and is consistent with many studies showing that increased host abundance increases specialist pathogen infection (Civitello *et al*. 2013; Parker *et al*. 2015; Halliday *et al*. 2020; Liu *et al*. 2023, but see Schmidt et al., 2020). The strong importance of dilution effects for fungal pathogens is likely due to them being host specific and spread at a local scale between susceptible neighbours (e.g., by wind, water droplets). Surprisingly, however, these dilution effects did not translate into lower pathogen damage in more diverse plots (Rottstock *et al*. 2014), which is consistent with previous results from this experiment (Cappelli et a. 2020). The lack of a diversity effect could be because resource concentration effects are counterbalanced by the spillover of more generalist pathogens between plant species in diverse plots, e.g., spillovers between highly infected fast species. Additionally, the strength of host concentration effects could vary between fast and slow species. Slow species showed stronger resource concentration effects and suffered more rust infection when growing in patches surrounded by conspecifics. As well defended species are expected to have more specialist consumers (Winde & Wittstock 2011; Bruce 2014), they might show stronger host concentration effects. The fact that the PaNDiv experiment contains diverse plots with only fast species (where dilution effects did not operate) may have reduced the strength of diversity effects, compared to other experiments (Rottstock *et al*. 2014). Overall, these findings show that dilution effects at the species level do not necessarily lead to negative diversity-disease relationships at the community level and they highlight the importance of community context in driving diversity-disease relationships.

Shifts in conspecific density were hypothesised to have differential effects on insect herbivore guilds, and as expected, chewing herbivory increased with decreasing conspecific density, while sucking herbivory did not. Sucking herbivores are expected to be predominantly specialists, and whilst we do see a positive effect of conspecific density on species level sucking herbivory, it was driven by fast growing species. This is the opposite pattern to that observed for rust pathogens and could suggest that slow species are so well defended that they do not suffer high sucking herbivore attack even at high conspecific density. The increase in chewing herbivory with increasing plant species richness, and the decrease in damage with increasing conspecific density, would be expected if chewing herbivores are generalists, which benefit from diet mixing in diverse plant communities (Bernays *et al*. 1994). Consumer guilds also interacted strongly. In control plots, chewing herbivory decreased with increasing conspecific density, however this relationship was reversed under fungicide application. As discussed previously, pathogen infection increased significantly with increasing conspecific density, suggesting the reduction in chewing damage on species growing in control plots with high conspecific density could be driven by chewing consumers avoiding plants heavily infected with pathogens (Jallow *et al*. 2004). These results highlight the importance of investigating individual effects of plant community shifts on different herbivore guilds, but also interactions between the various consumer guilds that simultaneously attack plants in natural communities (Conrath *et al*. 2002).

We disentangled the indirect mechanisms by which nitrogen enrichment affects consumer damage at different organisational levels and found strong support for the growth-defence trade-off and resource concentration hypotheses. However, growth-defence effects frequently emerged from interactions between species within a community (associational effects) and growth-defence and resource concentration effects often interacted with each other. It is therefore not straightforward to transfer patterns from the species to the community level, as effects could be amplified when scaling from species to communities, e.g., growth-defence effects for rust pathogens, while others were lost at the community level, e.g., a positive effect of conspecific density on pathogen infection at the species level did not result in a reduction in infection with increased diversity at the community level. We therefore need to consider interactions between plant species within communities if we are to determine how effects of diversity and functional trait composition affect consumer groups at different levels.

## Acknowledgements

We are grateful to the whole PaNDiv team, especially Hugo Vincent and Mervi Laitinen and many helpers, for maintaining the experiment and without whom the PaNDiv Experiment would not be possible. We also thank Seraina Cappelli and Noémie Pichon for their role in setting up the PaNDiv Experiment. Several people helped with collecting the plant cover data, in particular: Cindy Bucher, Tala Bürki, Nadia Maaroufi, Anja Michel, Barryette, Oberholzer, Noémie Pichon, Valentin Pulver, Fabian Heussler and Lia Zehnder. The project was funded by the Swiss National Science Foundation (Award 310030_185260).

## References

1. Alexander, H.M. & Jeanne, D.M. (2000). Seedling Disease in an Annual Legume: Consequences for Seedling Mortality, Plant Size, and Population Seed Production. Oecologia, 122, 346–353.

2. Ali, J.G. & Agrawal, A.A. (2012). Specialist versus generalist insect herbivores and plant defense. Spec. Issue Specif. Plant–enemy Interact., 17, 293–302.

3. Allan, E., van Ruijven, J. & Crawley, M.J. (2010). Foliar fungal pathogens and grassland biodiversity. Ecology, 91, 2572–2582.

4. Alm Bergvall, U., Rautio, P., Kesti, K., Tuomi, J. & Leimar, O. (2006). Associational effects of plant defences in relation to within- and between-patch food choice by a mammalian herbivore: neighbour contrast susceptibility and defence. Oecologia, 147, 253–260.

5. Atsatt, P. & O’Dowd, D. (1976). Plant defense guilds. Science, 193, 24–29.

6. Barbosa, P., Hines, J., Kaplan, I., Martinson, H., Szczepaniec, A. & Szendrei, Z. (2009). Associational resistance and associational susceptibility: Having right or wrong neighbors. *Annu. Rev. Ecol. Evol. Syst.*, Annual review of ecology evolution and systematics, 40, 1–20.

7. Behmer, S.T. & Joern, A. (2008). Coexisting generalist herbivores occupy unique nutritional feeding niches. Proc. Natl. Acad. Sci. U. S. A., 105, 1977–1982.

8. Bernays, E.A., Bright, K.L., Gonzalez, N. & Angel, J. (1994). Dietary Mixing in a Generalist Herbivore: Tests of Two Hypotheses. Ecology, 75, 1997–2006.

9. Berner, D., Blanckenhorn, W. & Korner, C. (2005). Grasshoppers cope with low host plant quality by compensatory feeding and food selection: N limitation challenged. Oikos Cph. Den., 111, 525–533.

10. Blüthgen, N., Dormann, C.F., Prati, D., Klaus, V.H., Kleinebecker, T., Hölzel, N., et al. (2012). A quantitative index of land-use intensity in grasslands: Integrating mowing, grazing and fertilization. Basic Appl. Ecol., 13, 207–220.

11. Borer, E.T., Harpole, W.S., Adler, P.B., Lind, E.M., Orrock, J.L., Seabloom, E.W., et al. (2014). Finding generality in ecology: a model for globally distributed experiments. Methods Ecol. Evol., 5, 65–73.

12. Brooks, M.E., Kristensen, K., van Benthem, K.J., Magnusson, A., Berg, C.W., Nielsen, A., et al. (2017). glmmTMB balances speed and flexibility among packages for zero-inflated generalized linear mixed modeling. R J., 9, 378–400.

13. Bruce, T.J.A. (2014). Glucosinolates in oilseed rape: secondary metabolites that influence interactions with herbivores and their natural enemies. Ann. Appl. Biol., 164, 348– 353.

14. Burdon, J.J., Thrall, P.H., Ericson, & Lars. (2006). The Current and Future Dynamics of Disease in Plant Communities. Annu. Rev. Phytopathol., 44, 19–39.

15. Cappelli, S.L., Pichon, N.A., Kempel, A. & Allan, E. (2020). Sick plants in grassland communities: a growth-defense trade-off is the main driver of fungal pathogen abundance. Ecol. Lett., 23, 1349–1359.

16. Champagne, E., Tremblay, J.-P. & Cote, S.D. (2016). Spatial extent of neighboring plants influences the strength of associational effects on mammal herbivory. Ecosphere Wash. DC, 7.

17. Civitello, D.J., Pearsall, S., Duffy, M.A. & Hall, S.R. (2013). Parasite consumption and host interference can inhibit disease spread in dense populations. Ecol. Lett., 16, 626–634.

18. Coley, P., Bryant, J. & Chapin, F. (1985). Resource availability and plant antiherbivore defense. Science, 230, 895–899.

19. Conrath, U., Pieterse, C.M.J. & Mauch-Mani, B. (2002). Priming in plant–pathogen interactions. Trends Plant Sci., 7, 210–216.

20. Díaz, S., Kattge, J., Cornelissen, J.H.C., Wright, I.J., Lavorel, S., Dray, S., et al. (2016). The global spectrum of plant form and function. Nature, 529, 167–171.

21. Dordas, C. (2008). Role of nutrients in controlling plant diseases in sustainable agriculture. A review. Agron. Sustain. Dev., 28, 33–46.

22. Dunn, P.K. (2021). Package‘tweedie’: Evaluation of Tweedie Exponential Family Models.

23. Ebeling, A., Strauss, A.T., Adler, P.B., Arnillas, C.A., Barrio, I.C., Biederman, L.A., et al. (2021). Nutrient enrichment increases invertebrate herbivory and pathogen damage in grasslands. J. Ecol.

24. Eberl, F., de Bobadilla, M.F., Reichelt, M., Hammerbacher, A., Gershenzon, J. & Unsicker, S.B. (2020). Herbivory meets fungivory: insect herbivores feed on plant pathogenic fungi for their own benefit. Ecol. Lett.

25. Eskelinen, A., Harpole, W.S., Jessen, M.-T., Virtanen, R. & Hautier, Y. (2022). Light competition drives herbivore and nutrient effects on plant diversity. Nature, 611, 301– 305.

26. Felton, G.W. & Korth, K.L. (2000). Trade-offs between pathogen and herbivore resistance. Curr. Opin. Plant Biol., 3, 309–314.

27. Fenner, M., Hanley, M. & Lawrence, R. (1999). Comparison of seedling and adult palatability in annual and perennial plants. Funct. Ecol., 13, 546–551.

28. Forister, M.L., Novotny, V., Panorska, A.K., Baje, L., Basset, Y., Butterill, P.T., et al. (2015). The global distribution of diet breadth in insect herbivores. Proc. Natl. Acad. Sci. U. S. A., 112, 442–447.

29. Garnier, E., Shipley, B., Roumet, C. & Laurent, G. (2001). A standardized protocol for the determination of specific leaf area and leaf dry matter content. Funct. Ecol., 15, 688– 695.

30. Halliday, F.W., Heckman, R.W., Wilfahrt, P.A. & Mitchell, C.E. (2017). A multivariate test of disease risk reveals conditions leading to disease amplification. Proc. R. Soc. B Biol. Sci., 284, 20171340.

31. Halliday, F.W., Heckman, R.W., Wilfahrt, P.A. & Mitchell, C.E. (2019). Past is prologue: host community assembly and the risk of infectious disease over time. Ecol. Lett., 22, 138–148.

32. Halliday, F.W., Rohr, J.R. & Laine, A.-L. (2020). Biodiversity loss underlies the dilution effect of biodiversity. Ecol. Lett., 23, 1611–1622.

33. Hanley, M.E., Fenner, M. & Edwards, P.J. (1995). The Effect of Seedling Age on the Likelihood of Herbivory by the Slug Deroceras reticulatum. Funct. Ecol., 9, 754–759.

34. Hartig, F. & Lohse, L. (2021). Package ‘DHARMa’: Residual Diagnostics for Hierarchical (Multi-Level / Mixed) Regression Models.

35. Hautier, Y., Niklaus, P.A. & Hector, A. (2009). Competition for light causes plant biodiversity loss after eutrophication. Science, 324, 636–638.

36. Herms, D. & Mattson, W. (1992). The dilemma of plants - To grow or defend. Q. Rev. Biol., 67, 283–335.

37. Hjalten, J., Danell, K. & Lundberg, P. (1993). Herbivore avoidance by association - Vole and hare utilization of woody-plants. Oikos Cph. Den., 68, 125–131.

38. Huber, D. & Watson, R. (1974). Nitrogen form and plant disease. Annu. Rev. Phytopathol., 12, 139–165.

39. Jallow, M.F.A., Dugassa-Gobena, D. & Vidal, S. (2004). Indirect interaction between an unspecialized endophytic fungus and a polyphagous moth. Basic Appl. Ecol., 5, 183– 191.

40. van Klink, R., Bowler, D.E., Gongalsky, K.B., Swengel, A.B., Gentile, A. & Chase, J.M. (2020). Meta-analysis reveals declines in terrestrial but increases in freshwater insect abundances. Science, 368, 417+.

41. Krieger, R.I., Feeny, P.P., & Christopher F. Wilkinson. (1971). Detoxication enzymes in the guts of caterpillars: An evolutionary answer to plant defenses? Science, 172, 579–581.

42. La Pierre, K.J. & Smith, M.D. (2016). Soil nutrient additions increase invertebrate herbivore abundances, but not herbivory, across three grassland systems. Oecologia, 180, 485– 497.

43. Lavorel, S. & Grigulis, K. (2012). How fundamental plant functional trait relationships scale-up to trade-offs and synergies in ecosystem services. J. Ecol., 100, 128–140.

44. Lind, E.M., Borer, E., Seabloom, E., Adler, P., Bakker, J.D., Blumenthal, D.M., et al. (2013). Life-history constraints in grassland plant species: a growth-defence trade-off is the norm. Ecol. Lett., 16, 513–521.

45. Liu, X., Lyu, S., Sun, D., Bradshaw, C.J.A. & Zhou, S. (2017). Species decline under nitrogen fertilization increases community-level competence of fungal diseases. Proc. R. Soc. B-Biol. Sci., 284.

46. Liu, X., Lyu, S., Zhou, S. & Bradshaw, C.J.A. (2016). Warming and fertilization alter the dilution effect of host diversity on disease severity. Ecology, 97, 1680–1689.

47. Liu, X., Xiao, Y., Lin, Z., Wang, X., Hu, K., Liu, M., et al. (2023). Spatial scale-dependent dilution effects of biodiversity on plant diseases in grasslands. Ecology, 104, e3944.

48. Lopes, T., Hatt, S., Xu, Q., Chen, J., Liu, Y. & Francis, F. (2016). Wheat (Triticum aestivum L.)-based intercropping systems for biological pest control. PEST Manag. Sci., 72, 2193–2202.

49. Loranger, H., Weisser, W.W., Ebeling, A., Eggers, T., De Luca, E., Loranger, J., et al. (2014). Invertebrate herbivory increases along an experimental gradient of grassland plant diversity. Oecologia, 174, 183–193.

50. Lüdecke, D. (2018). ggeffects: Tidy Data Frames of Marginal Effects from Regression Models. ggeffects.

51. Lüdecke, D. (2021). sjPlot: Data Visualization for Statistics in Social Science. sjPlot.

52. Mitchell, C.E., Reich, P.B., Tilman, D. & Groth, J.V. (2003). Effects of elevated CO2, nitrogen deposition, and decreased species diversity on foliar fungal plant disease. Glob. Change Biol., 9, 438–451.

53. Mordecai, E.A. (2011). Pathogen impacts on plant communities: unifying theory, concepts, and empirical work. Ecol. Monogr., 81, 429–441.

54. Mur, L.A.J., Simpson, C., Kumari, A., Gupta, A.K. & Gupta, K.J. (2017). Moving nitrogen to the centre of plant defence against pathogens. Ann. Bot., 119, 703–709.

55. Parker, I.M., Saunders, M., Bontrager, M., Weitz, A.P., Hendricks, R., Magarey, R., et al. (2015). Phylogenetic structure and host abundance drive disease pressure in communities. Nature, 520, 542–544.

56. Pfisterer, A.B., Diemer, M. & Schmid, B. (2003). Dietary shift and lowered biomass gain of a generalist herbivore in species-poor experimental plant communities. Oecologia, 135, 234–241.

57. Pichon, N.A., Cappelli, S.L., Soliveres, S., Hoelzel, N., Klaus, V.H., Kleinebecker, T., et al. (2020). Decomposition disentangled: A test of the multiple mechanisms by which nitrogen enrichment alters litter decomposition. Funct. Ecol., 34, 1485–1496.

58. Power, A.G. & Mitchell, C.E. (2004). Pathogen Spillover in Disease Epidemics. Am. Nat., 164, S79–S89.

59. R Core Team. (2021). R: A Language and Environment for Statistical Computing.

60. Root, R. (1973). Organization of a plant-arthropod association in simple and diverse habitats - fauna of collards (brassica-oleracea). Ecol. Monogr., 43, 95–120.

61. Rottstock, T., Joshi, J., Kummer, V. & Fischer, M. (2014). Higher plant diversity promotes higher diversity of fungal pathogens, while it decreases pathogen infection per plant. Ecology, 95, 1907–1917.

62. Sanson, G., Read, J., Aranwela, N., Clissold, F. & Peeters, P. (2001). Measurement of leaf biomechanical properties in studies of herbivory: Opportunities, problems and procedures. Austral Ecol., 26, 535–546.

63. Schmid, B., Hector, A., Huston, M., Inchausti, P., Nijs, I. & Leadley, P. (2002). The design and analysis of biodiversity experiments, 61–78.

64. Schmidt, R., Auge, H., Deising, H.B., Hensen, I., Mangan, S.A., Schaedler, M., et al. (2020). Abundance, origin, and phylogeny of plants do not predict community-level patterns of pathogen diversity and infection. Ecol. Evol., 10, 5506–5516.

65. Slansky, F. & Feeny, P. (1977). Stabilization of rate of nitrogen accumulation by larvae of cabbage butterfly on wild and cultivated food plants. Ecol. Monogr., 47, 209–228.

66. Stout, M., Thaler, J. & Thomma, B. (2006). Plant-mediated interactions between pathogenic microorganisms and herbivorous arthropods. *Annu. Rev. Entomol.*, Annual review of entomology, 51, 663–689.

67. Strengbom, J. & Reich, P.B. (2006). Elevated [CO2] and increased N supply reduce leaf disease and related photosynthetic impacts on Solidago rigida. Oecologia, 149, 519– 525.

68. Suding, K.N., Collins, S.L., Gough, L., Clark, C., Cleland, E.E., Gross, K.L., et al. (2005). Functional- and abundance-based mechanisms explain diversity loss due to N fertilization. Proc. Natl. Acad. Sci., 102, 4387–4392.

69. Tahvanainen, J.O. & Root, R.B. (1972). The Influence of Vegetational Diversity on the Population Ecology of a Specialized Herbivore, Phyllotreta cruciferae (Coleoptera: Chrysomelidae). Oecologia, 10, 321–346.

70. Thaler, J.S., Humphrey, P.T. & Whiteman, N.K. (2012). Evolution of jasmonate and salicylate signal crosstalk. Spec. Issue Specif. Plant–enemy Interact., 17, 260–270.

71. Thomas, C.D. (1986). Butterfly larvae reduce host plant survival in vicinity of alternative host species. Oecologia, 70, 113–117.

72. Throop, H. & Lerdau, M. (2004). Effects of nitrogen deposition on insect herbivory: Implications for community and ecosystem processes. Ecosyst. N. Y. N, 7, 109–133.

73. Wickham, H. (2016). ggplot2: Elegant Graphics for Data Analysis. ggplot2.

74. Winde, I. & Wittstock, U. (2011). Insect herbivore counteradaptations to the plant glucosinolate–myrosinase system. Plant-Insect Interact., 72, 1566–1575.

75. Wright, I.J., Reich, P.B., Westoby, M., Ackerly, D.D., Baruch, Z., Bongers, F., et al. (2004). The worldwide leaf economics spectrum. Nature, 428, 821–827.

