## Supplementary material for "Disentangling the mechanisms by which nitrogen enrichment affects consumer damage at different organisational levels"

*Table S1: PaNDiv experimental species categorised into functional group and growth form. Species in bold are species where consumer damage was measured on the phytometers.*

| PaNDiv experimental species | Functional group | Growth form |
|---|---|---|
| *Lolium perenne* | | |
| *Holcus lanatus* | Fast growing | |
| *Dactylis glomerata* | | |
| *Poa trivialis* | | Grasses |
| *Helictotrichon pubescens* | | |
| *Festuca rubra* | Slow growing | |
| *Bromus erectus* | | |
| *Anthoxantum odoratum* | | |
| *Crepis biennis* | | |
| *Taraxacum officinale* | | |
| *Gallium album* | Fast growing | |
| *Anthriscus sylvestris* | | |
| *Heracleum sphondylium* | | Herbs |
| *Rumex acetosa* | | |
| *Centaurea jacea* | | |
| *Plantago media* | | |
| *Salvia pratensis* | Slow growing | |
| *Daucus carota* | | |
| *Prunella grandiflora* | | |
| *Achillea millefolium* | | |

*Table S2: Model output from glmmTMB models for insect herbivore damage at the individual and species level. Individual level (40cm⌀), consumer damage measured as the percentage of leaves damaged on the focal individual; Species level (3m$^2$), insect herbivory measured as mean percentage of leaves damaged/plant/species/plot, and pathogen infection measured as the proportion of plants infected/species/plot.*

| | **Chewing: Population** | | | **Chewing: Individual** | | | **Sucking: Population** | | | **Sucking: Individual** | | |
|---|---|---|---|---|---|---|---|---|---|---|---|---|
| *Predictors* | *Estimates* | *std. Error* | *p* | *Estimates* | *std. Error* | *p* | *Estimates* | *std. Error* | *p* | *Estimates* | *std. Error* | *p* |
| Intercept | 0.11 (0.04 – 0.30) | 0.06 | **<0.001** | 0.09 (0.06 – 0.16) | 0.03 | **<0.001** | 0.43 (0.26 – 0.71) | 0.11 | **0.001** | 0.12 (0.06 – 0.23) | 0.04 | **<0.001** |
| Nitogen | 0.64 (0.36 – 1.13) | 0.19 | 0.127 | 1.56 (1.19 – 2.06) | 0.22 | **0.001** | 0.90 (0.71 – 1.14) | 0.11 | 0.373 | 0.85 (0.61 – 1.19) | 0.15 | 0.342 |
| Fungicide | 0.76 (0.44 – 1.33) | 0.22 | 0.342 | 0.99 (0.75 – 1.31) | 0.14 | 0.940 | 0.92 (0.73 – 1.17) | 0.11 | 0.510 | 1.01 (0.74 – 1.38) | 0.16 | 0.928 |
| Conspecific density | 0.83 (0.63 – 1.09) | 0.12 | 0.178 | 0.79 (0.63 – 0.99) | 0.09 | **0.037** | 1.07 (0.96 – 1.20) | 0.06 | 0.220 | 1.08 (0.87 – 1.34) | 0.12 | 0.478 |
| SLA CWM | 1.71 (1.16 – 2.53) | 0.34 | **0.007** | 1.09 (0.87 – 1.35) | 0.12 | 0.451 | 1.16 (0.97 – 1.40) | 0.11 | 0.109 | 0.96 (0.76 – 1.21) | 0.11 | 0.730 |
| Focal SLA | 0.68 (0.45 – 1.01) | 0.14 | 0.054 | 0.73 (0.54 – 0.98) | 0.11 | **0.035** | 1.03 (0.85 – 1.25) | 0.10 | 0.768 | 1.03 (0.77 – 1.36) | 0.15 | 0.853 |
| Nitrogen x SLA CWM | 0.94 (0.61 – 1.43) | 0.20 | 0.767 | 1.06 (0.87 – 1.30) | 0.11 | 0.551 | 0.87 (0.72 – 1.05) | 0.08 | 0.157 | 0.97 (0.77 – 1.22) | 0.11 | 0.790 |
| SLA CWM x Focal SLA | 0.76 (0.63 – 0.92) | 0.07 | **0.004** | 0.96 (0.88 – 1.06) | 0.05 | 0.455 | 0.94 (0.87 – 1.01) | 0.04 | 0.105 | 1.10 (1.00 – 1.22) | 0.06 | 0.059 |
| Focal SLA x Conspecific density | 0.85 (0.57 – 1.26) | 0.17 | 0.418 | 0.82 (0.59 – 1.15) | 0.14 | 0.245 | 1.21 (1.01 – 1.46) | 0.11 | **0.042** | 1.15 (0.79 – 1.66) | 0.22 | 0.470 |
| Fungicide x Conspecific density | 1.39 (1.05 – 1.84) | 0.20 | **0.023** | 1.04 (0.86 – 1.26) | 0.10 | 0.693 | 0.98 (0.87 – 1.10) | 0.06 | 0.699 | 0.94 (0.76 – 1.16) | 0.10 | 0.561 |
| Nitrogen x Conspecific density | 1.08 (0.82 – 1.43) | 0.15 | 0.567 | 1.02 (0.84 – 1.24) | 0.10 | 0.837 | 1.09 (0.97 – 1.22) | 0.06 | 0.138 | 0.84 (0.68 – 1.04) | 0.09 | 0.119 |
| Nitrogen x Fungicide | 1.50 (0.69 – 3.27) | 0.60 | 0.310 | 0.90 (0.62 – 1.29) | 0.17 | 0.553 | 1.08 (0.78 – 1.49) | 0.18 | 0.635 | 0.94 (0.59 – 1.48) | 0.22 | 0.779 |
| Nitrogen x Focal SLA | 0.98 (0.67 – 1.43) | 0.19 | 0.902 | 1.22 (0.98 – 1.53) | 0.14 | 0.073 | 0.95 (0.81 – 1.12) | 0.08 | 0.545 | 1.03 (0.84 – 1.27) | 0.11 | 0.784 |
| Fungicide x Focal SLA | 1.34 (0.91 – 1.96) | 0.26 | 0.136 | 1.07 (0.87 – 1.33) | 0.12 | 0.518 | 0.95 (0.81 – 1.11) | 0.08 | 0.520 | 0.89 (0.73 – 1.10) | 0.09 | 0.278 |
| Conspecific density x SLA CWM | 1.10 (0.84 – 1.45) | 0.15 | 0.481 | 1.06 (0.80 – 1.42) | 0.16 | 0.683 | 0.89 (0.78 – 1.02) | 0.06 | 0.099 | 0.89 (0.65 – 1.22) | 0.14 | 0.477 |
| **Random Effects** | | | | | | | | | | | | |
| $\sigma^2$ | 2.17 | | | 1.40 | | | 0.75 | | | 1.48 | | |
| $\tau_{00}$ | 0.10 $_{Block}$ | | | 0.00 $_{Block}$ | | | 0.04 $_{Block}$ | | | 0.10 $_{Block}$ | | |
| | 0.78 $_{Plot\_Nr}$ | | | 0.06 $_{Plot\_Nr:name}$ | | | 0.12 $_{Plot\_Nr}$ | | | 0.18 $_{Plot\_Nr:name}$ | | |
| | 0.03 $_{composition}$ | | | 0.06 $_{name}$ | | | 0.11 $_{composition}$ | | | 0.25 $_{name}$ | | |
| | 3.94 $_{Species}$ | | | 0.04 $_{composition}$ | | | 0.97 $_{Species}$ | | | 0.00 $_{composition}$ | | |
| | | | | 0.51 $_{Phy\_Species}$ | | | | | | 0.42 $_{Phy\_Species}$ | | |
| N | 4 $_{Block}$ | | | 4 $_{Block}$ | | | 4 $_{Block}$ | | | 4 $_{Block}$ | | |
| | 216 $_{Plot\_Nr}$ | | | 197 $_{Plot\_Nr}$ | | | 216 $_{Plot\_Nr}$ | | | 197 $_{Plot\_Nr}$ | | |
| | 54 $_{composition}$ | | | 12 $_{name}$ | | | 54 $_{composition}$ | | | 12 $_{name}$ | | |
| | 20 $_{Species}$ | | | 59 $_{composition}$ | | | 20 $_{Species}$ | | | 59 $_{composition}$ | | |
| | | | | 10 $_{Phy\_Species}$ | | | | | | 10 $_{Phy\_Species}$ | | |
| Observations | 1152 | | | 1624 | | | 1152 | | | 1624 | | |
| Marginal $R^2$ / Conditional $R^2$ | 0.045 / 0.705 | | | 0.071 / NA | | | 0.022 / 0.633 | | | 0.026 / NA | | |

*Table S3: Model output from glmmTMB model for community weighted mean (CWM) consumer damage. Community level (3m$^2$), consumer damage measured as community weighted mean (CWM) percentage of leaves damaged/plant/species/plot (insect herbivory), or as the proportion of plants infected/species/plot (pathogen infection). SLA= Specific leaf area.*

| | **Chewing** | | | **Sucking** | | | **Rust** | | | **Leafspot** | | |
|---|---|---|---|---|---|---|---|---|---|---|---|---|
| *Predictors* | *Estimates* | *std. Error* | *p* | *Estimates* | *std. Error* | *p* | *Estimates* | *std. Error* | *p* | *Estimates* | *std. Error* | *p* |
| Intercept | 0.13 (0.06 – 0.27) | 0.05 | **<0.001** | 0.67 (0.48 – 0.94) | 0.12 | **0.019** | 0.24 (0.16 – 0.38) | 0.05 | **<0.001** | 0.62 (0.39 – 0.86) | 0.12 | **<0.001** |
| Nitogen | 0.84 (0.58 – 1.21) | 0.16 | 0.342 | 0.93 (0.79 – 1.09) | 0.08 | 0.372 | 1.02 (0.77 – 1.34) | 0.14 | 0.916 | 0.00 (-0.05 – 0.06) | 0.03 | 0.900 |
| Fungicide | 1.37 (0.95 – 1.99) | 0.26 | 0.093 | 1.04 (0.89 – 1.23) | 0.09 | 0.612 | 0.24 (0.18 – 0.33) | 0.04 | **<0.001** | -0.03 (-0.08 – 0.03) | 0.03 | 0.299 |
| Species richness | 1.98 (1.12 – 3.50) | 0.57 | **0.018** | 1.03 (0.78 – 1.37) | 0.15 | 0.810 | 1.13 (0.80 – 1.58) | 0.20 | 0.488 | -0.04 (-0.09 – 0.01) | 0.03 | 0.147 |
| SLA CWM | 0.85 (0.51 – 1.41) | 0.22 | 0.527 | 1.11 (0.88 – 1.39) | 0.13 | 0.367 | 1.61 (1.18 – 2.18) | 0.25 | **0.002** | 0.01 (-0.05 – 0.06) | 0.03 | 0.811 |
| Nitrogen x SLA CWM | 1.50 (0.93 – 2.41) | 0.36 | 0.094 | 0.92 (0.75 – 1.13) | 0.09 | 0.417 | 0.77 (0.59 – 1.01) | 0.11 | 0.059 | -0.01 (-0.06 – 0.05) | 0.03 | 0.811 |
| Fungicide x SLA CWM | 1.08 (0.68 – 1.73) | 0.26 | 0.741 | 1.05 (0.85 – 1.29) | 0.11 | 0.652 | 1.54 (1.12 – 2.11) | 0.25 | **0.007** | 0.01 (-0.05 – 0.07) | 0.03 | 0.747 |
| Species richness x SLA CWM | 1.60 (1.07 – 2.40) | 0.33 | **0.023** | 1.14 (0.92 – 1.41) | 0.12 | 0.225 | 1.04 (0.79 – 1.37) | 0.15 | 0.769 | 0.03 (-0.01 – 0.08) | 0.02 | 0.151 |
| Nitrogen x Species richness | 0.93 (0.66 – 1.31) | 0.16 | 0.680 | 0.98 (0.83 – 1.16) | 0.09 | 0.833 | 1.01 (0.76 – 1.34) | 0.15 | 0.943 | -0.04 (-0.09 – 0.01) | 0.03 | 0.145 |
| Fungicide x Species richness | 0.99 (0.70 – 1.42) | 0.18 | 0.966 | 1.05 (0.88 – 1.26) | 0.10 | 0.561 | 0.93 (0.67 – 1.29) | 0.15 | 0.653 | 0.03 (-0.02 – 0.09) | 0.03 | 0.254 |
| **Random Effects** | | | | | | | | | | | | |
| $\sigma^2$ | 1.51 | | | 0.35 | | | 0.67 | | | 0.02 | | |
| $\tau_{00}$ | $0.14_{Block}$ | | | $0.04_{Block}$ | | | $0.08_{Block}$ | | | $0.05_{Block}$ | | |
| | $0.85_{Plot\_Nr}$ | | | $0.08_{Plot\_Nr}$ | | | $0.00_{Plot\_Nr}$ | | | $0.02_{Plot\_Nr}$ | | |
| | $2.88_{composition}$ | | | $0.73_{composition}$ | | | $0.82_{composition}$ | | | $0.00_{composition}$ | | |
| N | $4_{Block}$ | | | $4_{Block}$ | | | $4_{Block}$ | | | $4_{Block}$ | | |
| | $216_{Plot\_Nr}$ | | | $216_{Plot\_Nr}$ | | | $216_{Plot\_Nr}$ | | | $216_{Plot\_Nr}$ | | |
| | $54_{composition}$ | | | $54_{composition}$ | | | $54_{composition}$ | | | $54_{composition}$ | | |
| Observations | 216 | | | 216 | | | 216 | | | 216 | | |
| Marginal $R^2$ / Conditional $R^2$ | 0.094 / 0.698 | | | 0.010 / 0.690 | | | 0.558 / NA | | | 0.048 / 0.777 | | |

*Table S4: Model output from glmmTMB models for fungal pathogen damage at the Individual and Species level. Individual level (40cm⌀), consumer damage measured as the percentage of leaves damaged on the focal individual; Species level(3m$^2$), insect herbivory measured as mean percentage of leaves damaged/plant/species/plot, and pathogen infection measured as the proportion of plants infected/species/plot.*

| | **Rust: Population** | | | **Rust: Individual** | | | **Leafspot: Population** | | | **Leafspot: Individual** | | |
|---|---|---|---|---|---|---|---|---|---|---|---|---|
| *Predictors* | *Estimates* | *std. Error* | *p* | *Estimates* | *std. Error* | *p* | *Estimates* | *std. Error* | *p* | *Estimates* | *std. Error* | *p* |
| Intercept | 0.03<br>(0.01 – 0.12) | 0.02 | **<0.001** | 0.00<br>(0.00 – 0.02) | 0.00 | **<0.001** | 0.36<br>(0.19 – 0.68) | 0.12 | **0.002** | 0.30<br>(0.21 – 0.42) | 0.05 | **<0.001** |
| Nitogen | 1.05<br>(0.86 – 1.27) | 0.10 | 0.626 | 0.86<br>(0.40 – 1.85) | 0.34 | 0.700 | 0.94<br>(0.85 – 1.05) | 0.05 | 0.276 | 0.95<br>(0.82 – 1.11) | 0.07 | 0.532 |
| Fungicide | 0.27<br>(0.22 – 0.33) | 0.03 | **<0.001** | 0.22<br>(0.10 – 0.48) | 0.09 | **<0.001** | 1.00<br>(0.90 – 1.11) | 0.05 | 0.931 | 0.89<br>(0.76 – 1.03) | 0.07 | 0.111 |
| Conspecific density | 1.26<br>(1.12 – 1.42) | 0.08 | **<0.001** | 1.07<br>(0.54 – 2.10) | 0.37 | 0.848 | 1.13<br>(1.04 – 1.22) | 0.04 | **0.002** | 1.25<br>(1.12 – 1.40) | 0.07 | **<0.001** |
| SLA CWM | 1.11<br>(0.93 – 1.33) | 0.10 | 0.248 | 0.54<br>(0.25 – 1.17) | 0.21 | 0.119 | 0.98<br>(0.88 – 1.09) | 0.05 | 0.713 | 1.03<br>(0.91 – 1.17) | 0.07 | 0.596 |
| Focal SLA | 1.04<br>(0.81 – 1.34) | 0.13 | 0.766 | 1.84<br>(0.79 – 4.29) | 0.79 | 0.156 | 1.06<br>(0.93 – 1.22) | 0.07 | 0.377 | 1.03<br>(0.91 – 1.17) | 0.07 | 0.657 |
| Nitrogen x SLA CWM | 0.95<br>(0.79 – 1.16) | 0.09 | 0.636 | 1.23<br>(0.57 – 2.68) | 0.49 | 0.601 | 1.07<br>(0.95 – 1.20) | 0.06 | 0.278 | 0.98<br>(0.86 – 1.12) | 0.07 | 0.782 |
| SLA CWM x Focal SLA | 1.10<br>(1.01 – 1.20) | 0.05 | **0.029** | 1.10<br>(0.81 – 1.50) | 0.17 | 0.554 | 1.03<br>(0.98 – 1.08) | 0.03 | 0.307 | 1.03<br>(0.98 – 1.07) | 0.02 | 0.267 |
| Focal SLA x Conspecific density | 0.74<br>(0.59 – 0.92) | 0.08 | **0.006** | 1.07<br>(0.31 – 3.68) | 0.67 | 0.916 | 0.91<br>(0.79 – 1.03) | 0.06 | 0.137 | 0.96<br>(0.81 – 1.12) | 0.08 | 0.593 |
| Fungicide x Conspecific density | 1.12<br>(0.96 – 1.30) | 0.09 | 0.136 | 2.07<br>(1.02 – 4.24) | 0.76 | **0.045** | 0.95<br>(0.88 – 1.04) | 0.04 | 0.265 | 0.89<br>(0.80 – 1.00) | 0.05 | **0.047** |
| Nitrogen x Conspecific density | 0.92<br>(0.80 – 1.06) | 0.07 | 0.257 | 0.62<br>(0.26 – 1.47) | 0.27 | 0.281 | 1.04<br>(0.96 – 1.12) | 0.04 | 0.389 | 1.00<br>(0.89 – 1.12) | 0.06 | 0.980 |
| Nitrogen x Focal SLA | 1.05<br>(0.84 – 1.32) | 0.12 | 0.674 | 1.09<br>(0.58 – 2.06) | 0.35 | 0.786 | 0.96<br>(0.86 – 1.08) | 0.06 | 0.497 | 1.07<br>(0.97 – 1.17) | 0.05 | 0.169 |
| Conspecific density x SLA CWM | 1.14<br>(0.98 – 1.33) | 0.09 | 0.094 | 1.27<br>(0.47 – 3.43) | 0.64 | 0.641 | 1.02<br>(0.93 – 1.12) | 0.05 | 0.684 | 1.05<br>(0.91 – 1.20) | 0.07 | 0.509 |
| **Random Effects** | | | | | | | | | | | | |
| $\sigma^2$ | 2.38 | | | 5.44 | | | 0.49 | | | 0.52 | | |
| $\tau_{00}$ | $0.02_{\text{Block}}$<br>$0.01_{\text{Plot\_Nr}}$<br>$0.00_{\text{composition}}$<br>$6.75_{\text{Species}}$ | | | $0.24_{\text{Block}}$<br>$0.21_{\text{Plot\_Nr:name}}$<br>$2.69_{\text{name}}$<br>$0.00_{\text{composition}}$<br>$3.43_{\text{Phy\_Species}}$ | | | $0.37_{\text{Block}}$<br>$0.02_{\text{Plot\_Nr}}$<br>$0.00_{\text{composition}}$<br>$0.21_{\text{Species}}$ | | | $0.01_{\text{Block}}$<br>$0.15_{\text{Plot\_Nr:name}}$<br>$0.18_{\text{name}}$<br>$0.00_{\text{composition}}$<br>$0.06_{\text{Phy\_Species}}$ | | |
| N | $4_{\text{Block}}$<br>$216_{\text{Plot\_Nr}}$<br>$54_{\text{composition}}$<br>$20_{\text{Species}}$ | | | $4_{\text{Block}}$<br>$197_{\text{Plot\_Nr}}$<br>$12_{\text{name}}$<br>$59_{\text{composition}}$<br>$10_{\text{Phy\_Species}}$ | | | $4_{\text{Block}}$<br>$216_{\text{Plot\_Nr}}$<br>$54_{\text{composition}}$<br>$20_{\text{Species}}$ | | | $4_{\text{Block}}$<br>$197_{\text{Plot\_Nr}}$<br>$12_{\text{name}}$<br>$59_{\text{composition}}$<br>$10_{\text{Phy\_Species}}$ | | |
| Observations | 1152 | | | 1624 | | | 1152 | | | 1624 | | |
| Marginal $R^2$ / Conditional $R^2$ | 0.192 / NA | | | 0.224 / NA | | | 0.025 / 0.566 | | | 0.052 / 0.461 | | |

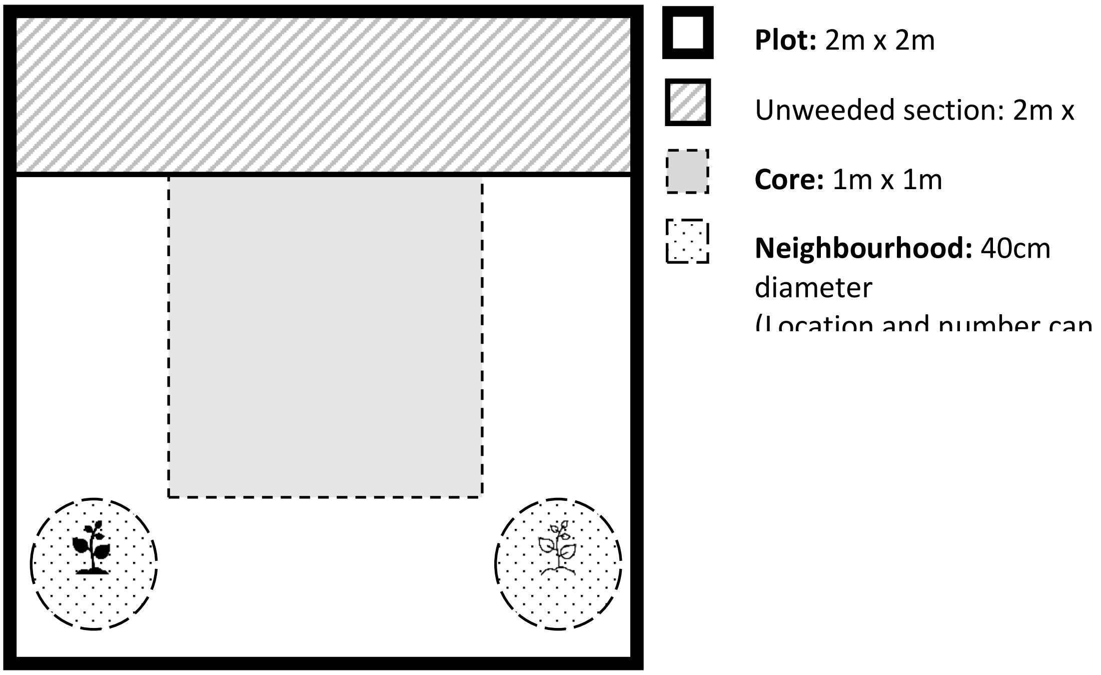


*Figure S1 : PaNDiv experimental plot layout.*

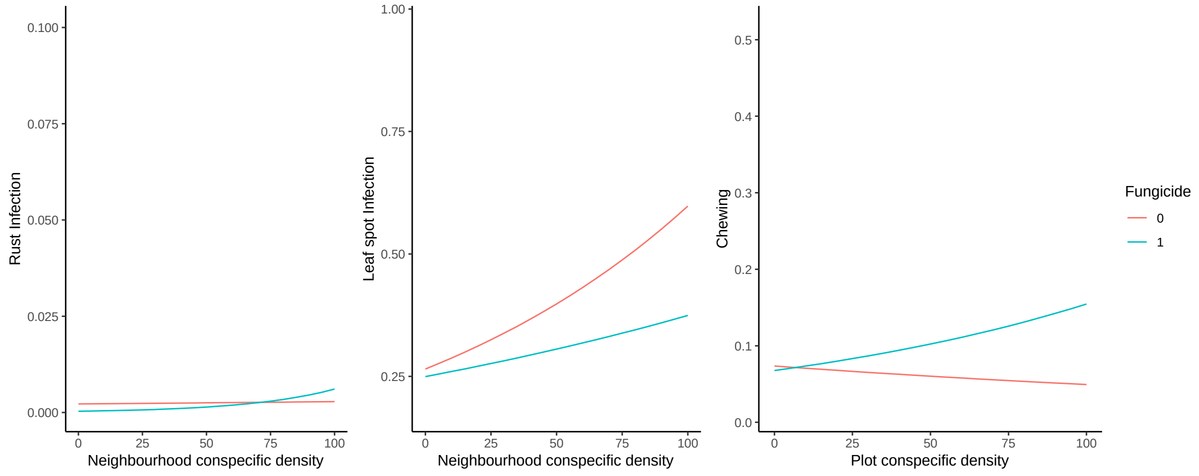


*Figure S2: Effect of the interaction between conspecific density and fungicide on consumer damage. Regression line with 95% confidence intervals, predicted from glmmTMB models are shown. Left and middle plots: Individual level= spatial scale 40cm⌀, consumer damage measured as the percentage of leaves damaged on the focal individual; Right plot: Species level = spatial scale 3m$^2$, insect herbivory measured as mean percentage of leaves damaged/plant/species/plot.*
